# Neutralizing antibody-dependent and -independent immune responses against SARS-CoV-2 in cynomolgus macaques

**DOI:** 10.1101/2020.08.18.256446

**Authors:** Hirohito Ishigaki, Misako Nakayama, Yoshinori Kitagawa, Cong Thanh Nguyen, Kaori Hayashi, Masanori Shiohara, Bin Gotoh, Yasushi Itoh

## Abstract

Severe acute respiratory syndrome coronavirus-2 (SARS-CoV-2) infectious disease (COVID-19) has been threatening the world because of severe symptoms and relatively high mortality. To develop vaccines and antiviral drugs for COVID-19, an animal model of SARS-CoV-2 infection is required to evaluate the efficacy of prophylactics and therapeutics *in vivo*. Therefore, we examined the pathogenicity of SARS-CoV-2 in cynomolgus macaques until 28 days after virus inoculation in the present study. Cynomolgus macaques showed body temperature rises after infection and X-ray radiographic viral pneumonia was observed in one of three macaques. However, none of the macaques showed life-threatening clinical signs of disease corresponding that approximately 80% of human patients did not show a critical disease in COVID-19. A neutralizing antibody against SARS-CoV-2 and T-lymphocytes that produced interferon (IFN)-γ and interleukin (IL)-2 specifically for SARS-CoV-2 N protein were detected on day 14 in the macaque that showed viral pneumonia. On the other hand, in the other macaques, in which a neutralizing antibody was not detected, T-lymphocytes that produced IFN-γ specifically for SARS-CoV-2 N protein increased on day 7 to day 14 prior to an increase in the number of T-lymphocytes that produced IL-2. These results suggest that not only a neutralizing antibody but also cellular immunity augmented by IFN-γ has a role in the elimination of SARS-CoV-2. Thus, because of the mild clinical signs of disease and low/no antibody responses against SARS-CoV-2 in two thirds of the macaques, cynomolgus macaques are appropriate to extrapolate human responses in vaccine and drug development.

**Author Summary:** Severe acute respiratory syndrome coronavirus-2 (SARS-CoV-2) infectious disease (COVID-19) has been threatening the world. To develop vaccines and antiviral drugs for COVID-19, an animal model of SARS-CoV-2 infection is required to evaluate their efficacy *in vivo*. Therefore, we examined the pathogenicity of SARS-CoV-2 in a non-human primate model until 28 days after virus inoculation. Cynomolgus macaques showed a fever after infection and X-ray radiographic viral pneumonia was observed in one of three macaques. However, none of the macaques showed life-threatening symptoms. A neutralizing antibody against SARS-CoV-2 and T-lymphocytes that produced interferon (IFN)-γ and interleukin (IL)-2 specifically for SARS-CoV-2 protein were detected on day 14 in the macaque that showed viral pneumonia. In the other macaques, in which a neutralizing antibody was not detected, T-lymphocytes that produced IFN-γ specifically for SARS-CoV-2 N protein increased on day 7 to day 14. These results suggest that not only a neutralizing antibody but also cellular immunity augmented by IFN-γ has a role in the elimination of SARS-CoV-2. Thus, because of the mild symptoms and low/no antibody responses against SARS-CoV-2 in two thirds of the macaques, cynomolgus macaques are appropriate to extrapolate human responses in vaccine and drug development.

## Introduction

Severe acute respiratory syndrome coronavirus-2 (SARS-CoV-2) infection (COVID-19) has been spreading around the world since late 2019 [1], and WHO declared a pandemic on March 11, 2020. Accumulating reports indicate varying degrees of illness including asymptomatic patients, patients with mild respiratory symptoms, and patients with acute respiratory distress syndrome (ARDS) requiring admission to the intensive care unit (ICU) [2–4]. In addition to the development of vaccines and antiviral drugs specific for SARS-CoV-2, determination of the pathogenicity in patients with severe clinical signs of disease and development of therapeutics for severe cases are urgent issues.

For the development of prophylactics and therapeutics for SARS-CoV-2 infection, not only *in vitro* studies but also *in vivo* studies are required for evaluation of their efficacy, especially estimation of *in vivo* efficacy, for which assessments are difficult in clinical trials. Therefore, animal models that show pathogenicity similar to that in humans are necessary for research and development of vaccines and antiviral drugs [5]. The results of several studies on the experimental infection of SARS-CoV-2 in animals have been reported. In a mouse model, SARS-CoV-2 propagated in the lungs of human angiotensin-converting enzyme 2 (ACE2) transgenic mice but not the lungs of wild-type mice, and the virus caused interstitial pneumonia in the ACE2 transgenic mice [6]. However, co-expression of human ACE2 and endogenous mouse ACE2 may change the disease progression to recapitulate COVID-19. Wild-type Syrian hamsters are sensitive to SARS-CoV-2, which propagated in the lungs to cause viral pneumonia, indicating a useful small animal model [7, 8]. However, since the viral pneumonia was resolved within 2 weeks in Syrian hamsters and antibodies that react to hamster molecules are not available in order to examine immune responses, another model is required to examine the pathogenicity of severe COVID-19. SARS-CoV-2 also propagated and caused lung inflammation and pneumonia in rhesus and cynomolgus macaques [9–13]. The pathogenicity in the macaques was examined until 21 days after virus infection in most of the studies except one study [13], in which the reason that the prolonged detection of viral genes in patients and virus-antigen specific T-lymphocyte responses were not revealed [14]. Therefore, in the present study, we observed cynomolgus macaques infected with SARS-CoV-2 for 4 weeks and examined T-lymphocyte responses specific for SARS-CoV-2 antigen peptides.

The macaque model, of which immune responses and metabolism resemble those of humans, is useful to extrapolate the efficacy of vaccines and antiviral drugs in humans against SARS-CoV-2. In our previous studies on influenza virus infection, various influenza viruses including pandemic and avian influenza viruses propagated in cynomolgus macaques that showed clinical signs of disease similar to human symptoms [15, 16]. In addition, we detected influenza viruses that were less sensitive to neuraminidase inhibitors in treated macaques, indicating a useful model for predicting the emergence of a drug-resistant virus [17, 18]. Therefore, we have used the cynomolgus macaque model to evaluate the efficacy of vaccines and antiviral drugs in influenza virus infection [19–23]. In the present study, we have expanded our experimental system to establish a SARS-CoV-2 infection model in cynomolgus macaques for preclinical studies.

We revealed the pathogenicity of SARS-CoV-2 in the cynomolgus macaques. SARS-CoV-2 propagated in respiratory tissues and caused body temperature rises in all of the macaques. However, viral pneumonia in X-ray radiographs was confirmed in one third of the macaques, in which a neutralizing antibody against SARS-CoV-2 in plasma was detected. We also found a thrombus in the lung of a macaque infected with SARS-CoV-2 as reported in human cases [24]. These results are similar to observations in human patients with COVID-19 [1]. Compared to influenza virus infection, the rate of detection of a neutralizing antibody was low in macaques infected with SARS-CoV-2 [19, 25]. In addition, interferon (IFN)-γ responses with delayed interleukin (IL)-2 responses might be related to virus elimination without a neutralizing antibody. Thus, cynomolgus macaques are an appropriate animal model of SARS-CoV-2 infection for the development of vaccines and antiviral drugs.

## Results

### SARS-CoV-2 propagation predominantly in the nasal and oral cavities of cynomolgus macaques

We inoculated the SARS-CoV-2 JPN/TY/WK-521/2020 (WK-521) strain [26] into the conjunctiva, nasal cavity, oral cavity, and trachea of cynomolgus macaques to examine the pathogenicity of the strain. We collected swab samples to examined virus propagation in the macaques. The virus was detected in swab samples of the conjunctiva, nasal cavity, oral cavity, and trachea of three macaques on day 1 (the next day after virus inoculation) (Table 1). The virus was detected in nasal and oral swab samples of two macaques, CE0202M and CE0324F, until day 7. No virus was detected in swab samples after day 10 or in homogenized tissue samples collected on day 28 at autopsy (listed in Table S1).

**Table 1.**
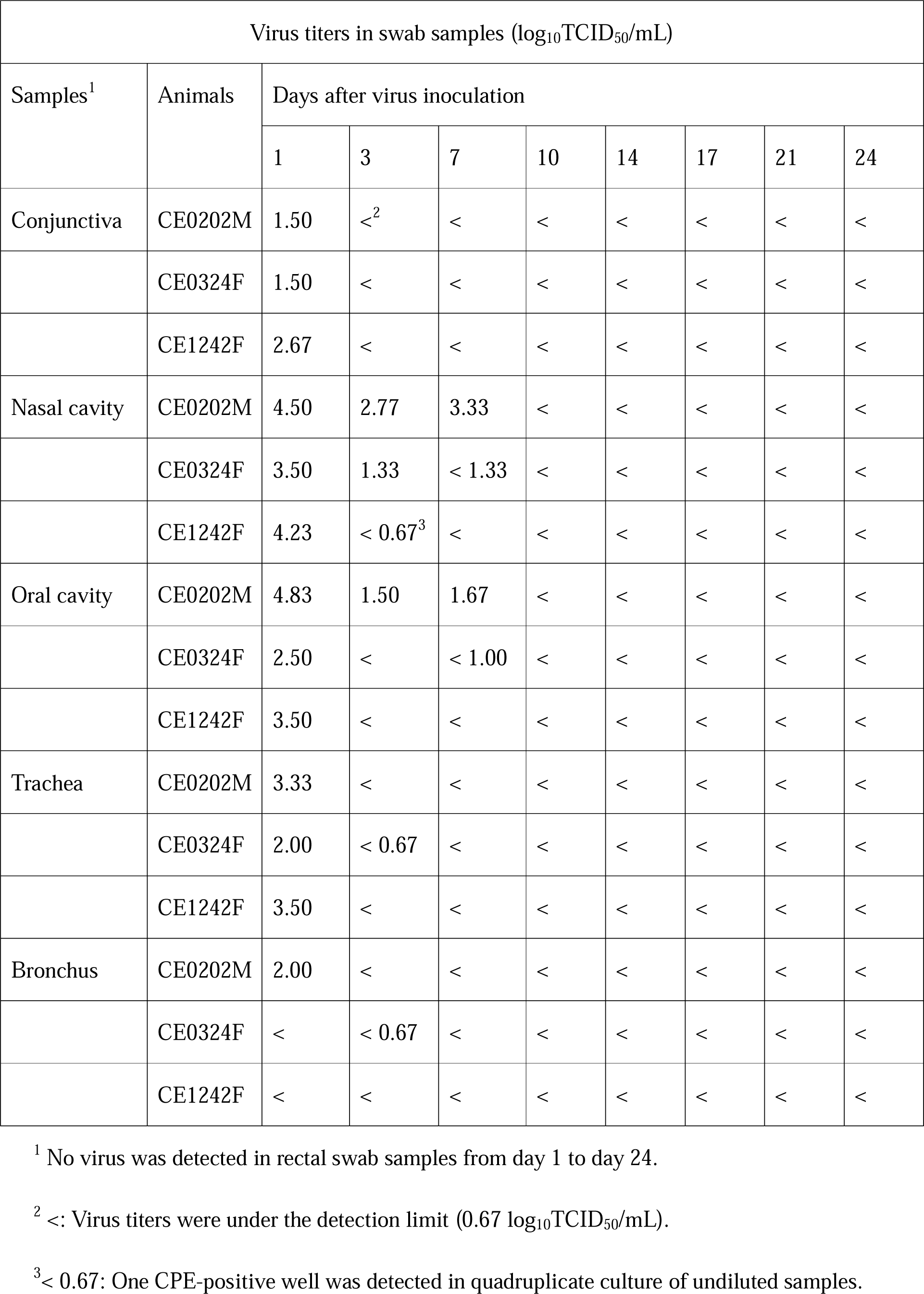
Virus titers in swab samples from cynomolgus macaques infected with SARS-CoV-2.

### Clinical signs of disease in macaques infected with SARS-CoV-2

Cynomolgus macaques showed clinical signs of disease after infection with SARS-CoV-2 WK-521. All of the macaques showed rises in body temperature after virus inoculation (day 1 to day 2) (Fig. 1). Body temperatures of two macaques (CE0324F and CE1242F) on day 1 were higher than 39.5oC, whereas CE0202M showed a rise in body temperature only at night from day 1 to day 2. Body weight of CE0324F decreased by 10% until day 28 due to loss of appetite (Fig. S1A). No macaques showed clinical scores that reached a humane endpoint (Table S2). Therefore, infection with WK-521 induced mild clinical signs of disease that were varied in the macaques.

**Fig. 1.**
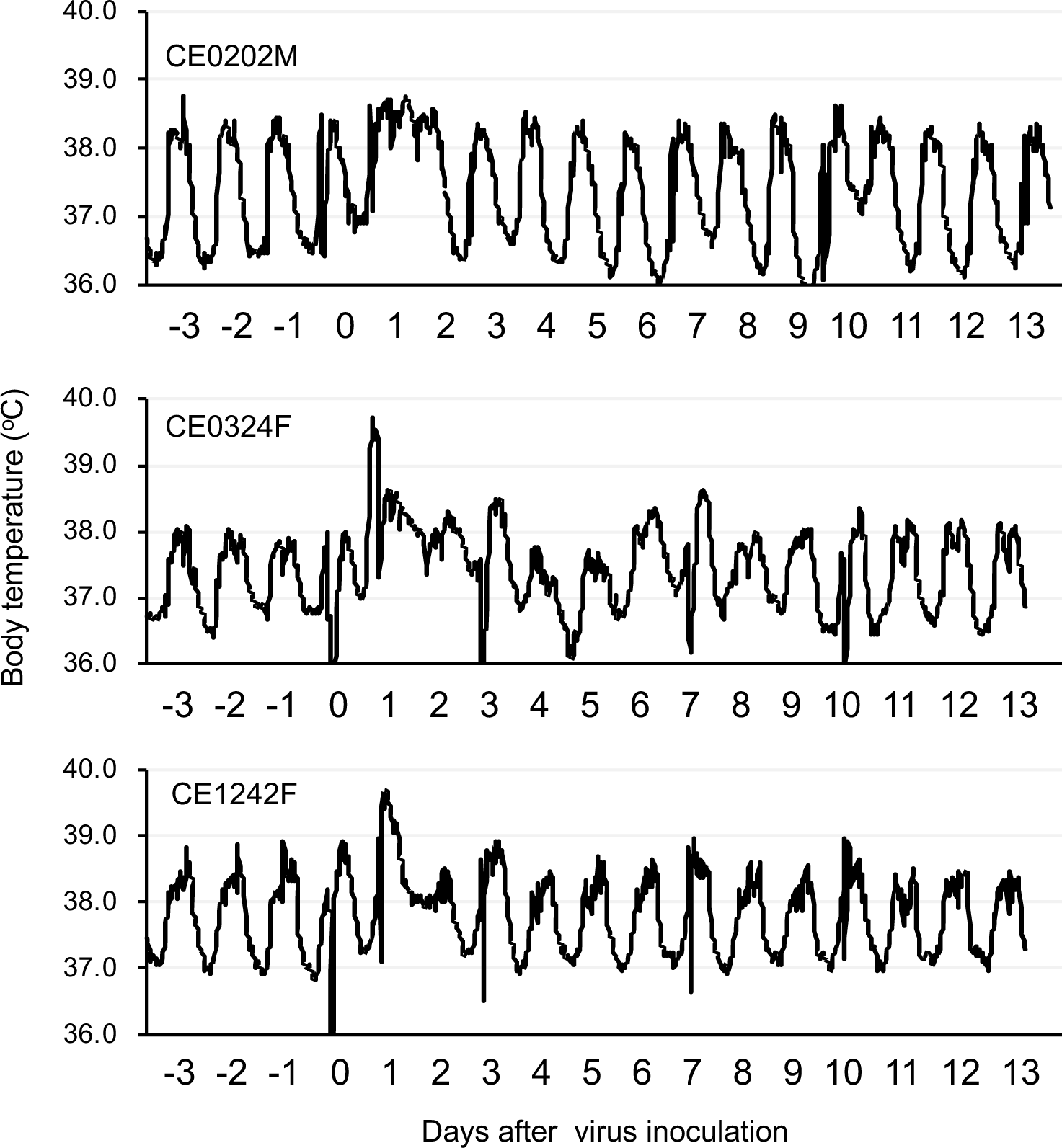
Body temperature change in cynomolgus macaques after SARS-CoV-2 infection. Cynomolgus macaques (n = 3) were inoculated with WK-521 on day 0. Body temperatures of the macaques were recorded using telemetry transmitters and a computer.

### Viral pneumonia in macaques infected with SARS-CoV-2

Viral pneumonia and/or bronchopneumonia was observed in cynomolgus macaques infected with SARS-CoV-2. Chest X-ray radiographs revealed a ground glass appearance in the lung, especially in the peripheral pulmonary areas, of CE0324F from day 3 to day 7 but not after day 10 (Fig. 2A–C). Saturation of percutaneous oxygen (SpO_2_) values in infected macaques at each sampling were higher than 90% during the study (Fig. S1B). At autopsy on day 28, gross lesions were observed on the surfaces of lungs including dark red and brown lesions in CE0324F, bright redness in CE0202M, and small red areas in CE1242F (Fig. 2D–F).

**Fig. 2.**
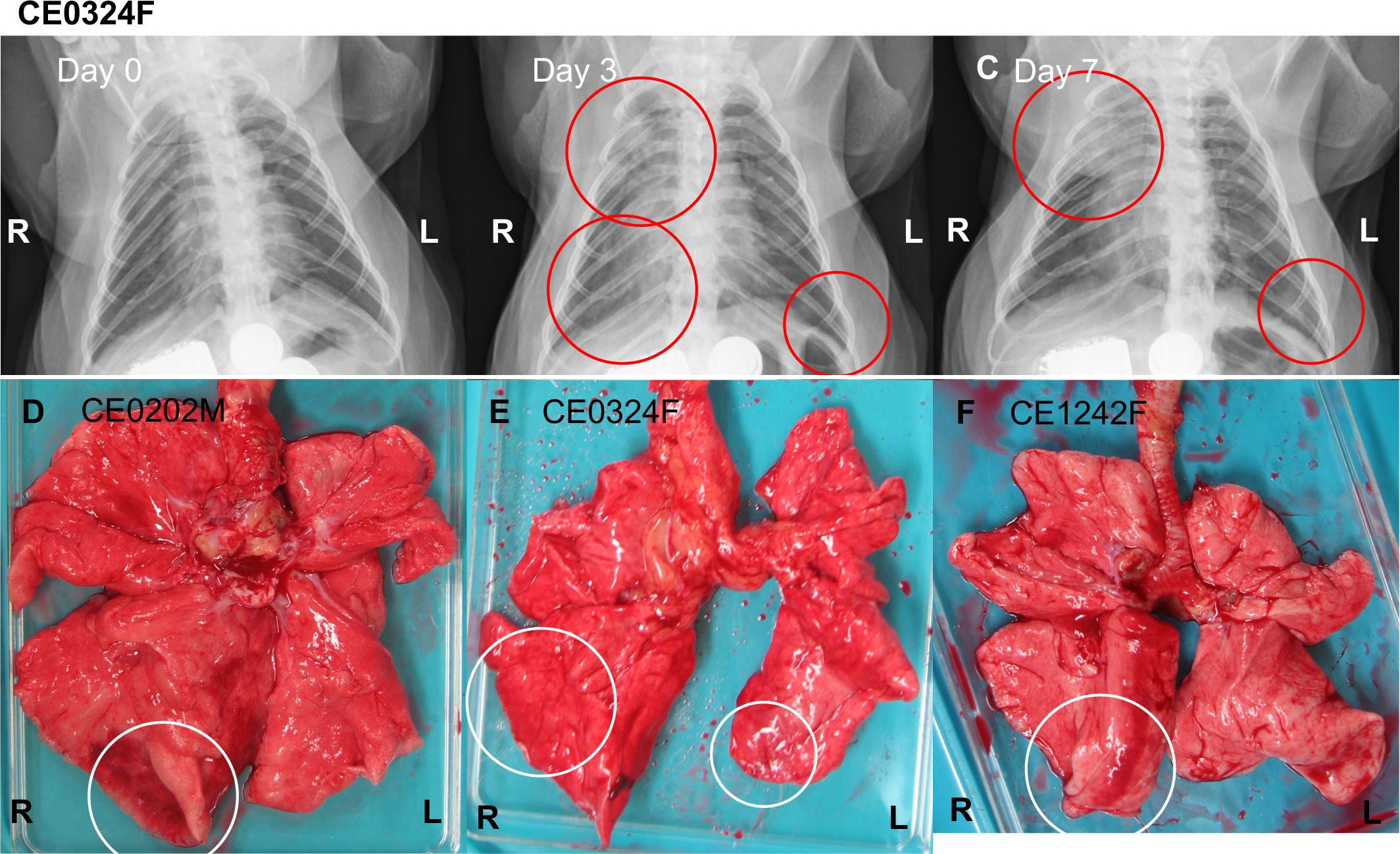
Viral pneumonia in cynomolgus macaques infected with SARS-CoV-2. (A-C) X-ray radiographs of CE0324F taken on day 0 before infection (A) and day 3 (B) and day 7 (C) after virus inoculation. Red circles indicate ground glass appearance (pneumonia). (D-F) Lung macroscopic appearance. All of the macaques infected with the virus were autopsied 28 days after virus inoculation. White circles indicate lesions with a bright red or dark red/brown color.

Histological pneumonia was observed in all of the macaques infected with WK-521. Thickened alveolar walls with infiltration of macrophages were observed in the lung with the macroscopic lesions in CE0324F, indicating a regenerative tissue (Fig. 3A). A larger number of bronchus-associated lymphoid tissues (BALT) were formed in the lung tissues of CE0324F (Fig. 3B) and CE1242F (Fig. S2A) than in those of CE0202M (Fig S2B). In CE0202M, congestion and a thrombus in the lung blood vessels were detected (Fig. 3D, E) and bronchopneumonia was detected in CE1242F (Fig. 3F). Lymphoid infiltration was detected in tissues other than respiratory organs. Focal lymphoid accumulation was seen in the submandibular gland of CE0324F (Fig. 3C). Thus, although no virus was detected on day 28, lymphoid responses continued until day 28 after virus inoculation.

**Fig. 3.**
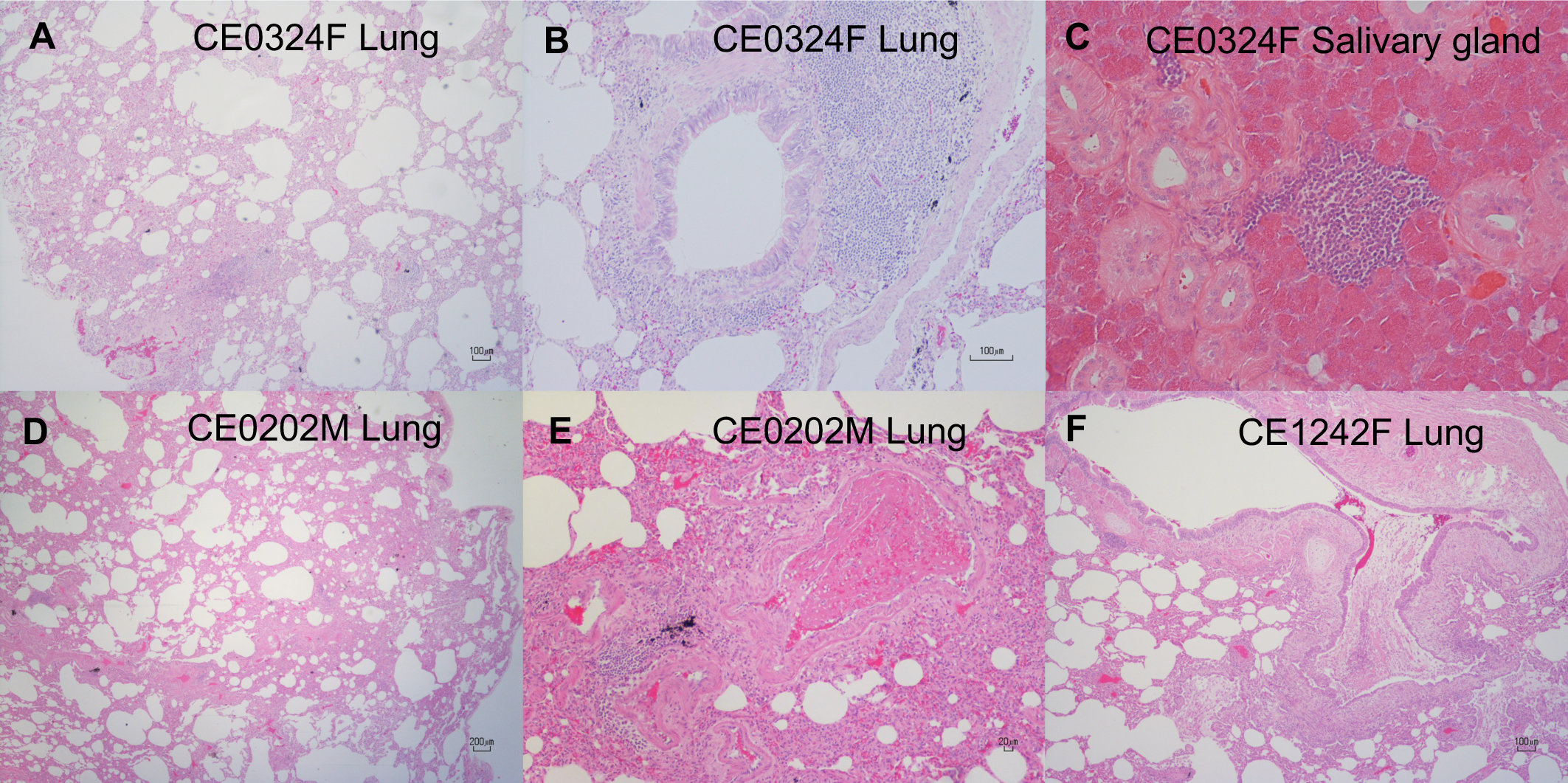
Inflammation in lungs and salivary glands of cynomolgus macaques infected with SARS-CoV-2. Lungs and salivary glands were collected on day 28 from macaques infected with SARS-CoV-2. HE-stained sections of lung tissues (A, B, D-F) and salivary glands (C). (A-C) CE0324F, (D, E) CE0202M, (F) CE1242F.

### Biochemical analysis of blood in macaques infected with SARS-CoV-2

Organ functions after infection with SARS-CoV-2 were biochemically analyzed. Levels of alanine aminotransferase (ALT) and alkaline phosphatase (ALP) in the plasma of CE0324F and CE1242F were increased after virus infection, but their levels including total bilirubin were within normal ranges, indicating minimal damage of the liver (Fig. 4A–C). No increase in the level of blood urea nitrogen (BUN) or creatinine was detected, indicating normal kidney function after virus infection (Fig. 4D, E). In CE0324F, the level of plasma amylase was temporally increased on day 1 (Fig. 4F).

**Fig. 4.**
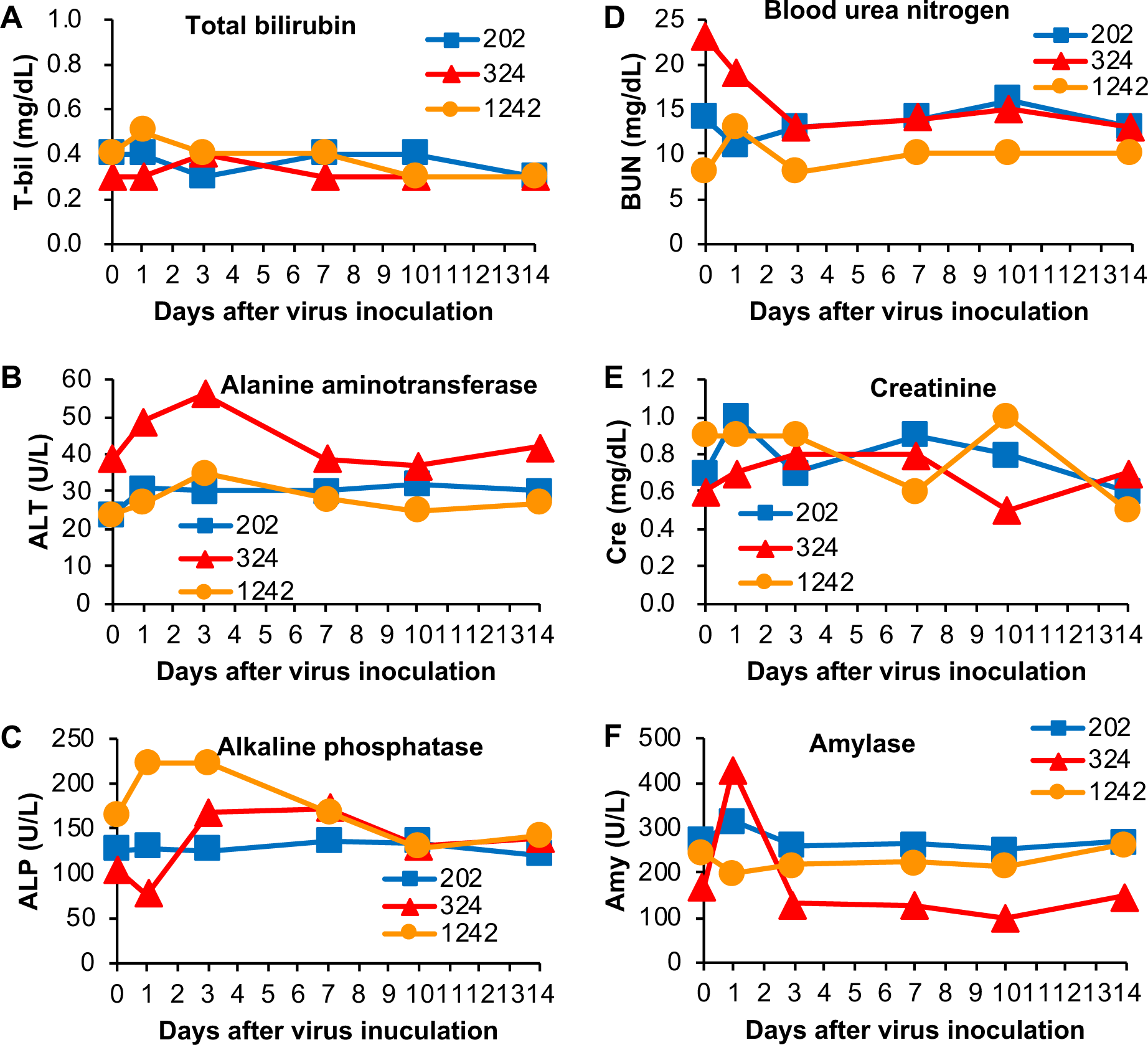
Blood biochemistry in cynomolgus macaques infected with SARS-CoV-2. Biochemical analysis was performed using plasma collected on the indicated days after virus inoculation: (A) total bilirubin, (B) alanine aminotransferase, (C) alkaline phosphatase, (D) blood urea nitrogen, (E) creatinine, and (F) amylase.

### Blood cell population in macaques infected with SARS-CoV-2

The blood cell population in macaques infected with SARS-CoV-2 was examined. The numbers of total white blood cells and granulocytes in CE0202M and CD1242F were increased temporally on day 1, but the number of total white blood cells in CE0324F was decreased on days 1 and 3 and it was increased after day 14 (Fig. 5A, B). The numbers of lymphocytes in the blood of CE0324F and CE1242F were decreased on day 1 and returned to the levels before infection on day 7 to day 10 (Fig. 5C), whereas the numbers of monocytes in CE0324F and CE1242F were increased on day 7 and day 3, respectively (Fig. 5D). The numbers of platelets in CE0324F and CE1242F were decreased on day 3. Thus, two of three macaques (CE0324F and CE1242F) showed apparent changes in the population of blood cells after SARS-CoV-2 infection.

**Fig. 5.**
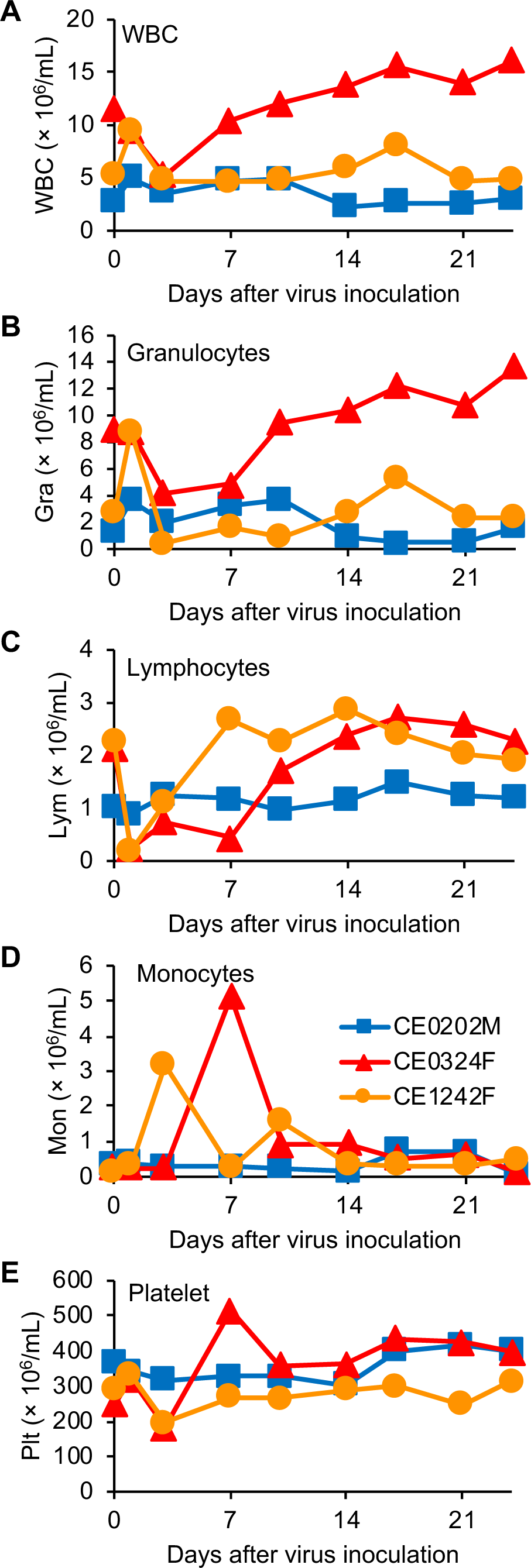
Blood cell counts in cynomolgus macaques infected with SARS-CoV-2. Peripheral blood cells were collected on the indicated days after virus inoculation. The concentrations of (A) total white blood cells, (B) granulocytes, (C) lymphocytes, (D) monocytes, and (E) platelets were determined.

### Immune responses in macaques infected with SARS-CoV-2

Immune responses against SARS-CoV-2 in the cynomolgus macaques were examined. CE0324F showed marked changes in plasma cytokine levels after infection (Fig. 6). The levels of the inflammatory cytokine interleukin-6 (IL-6) together with levels of monocyte chemotactic protein-1 (MCP-1) and IL-10 on day 1 were higher than those before infection. Levels of the Th2 cytokines IL-4 and IL-13, Th17 cytokine IL-17, and chemokines MIP-1α were increased 10 days after virus infection and the level of IL-8 was increased 21 days after virus infection, although the changes in IL-17 and MIP-1α were small. Levels of the Th1 cytokines interferon-γ (IFN-γ) and IL-12 were increased on day 1 after infection. IL-2 and TNF-α was detected in the plasma of CE0324F before infection (day 0). CE1242F showed low cytokine responses in IL-6, MCP-1, IL-10, IL-4, IL-13, MIP-1α, IFN-γ, and IL-12 compared with those in CE0324F. In CE0202M, levels of IL-8 and IL-12 on day 3 and levels of IL-13 from day 10 to day 17 were transiently increased, but no apparent changes in other cytokines were detected during the study.

**Fig. 6.**
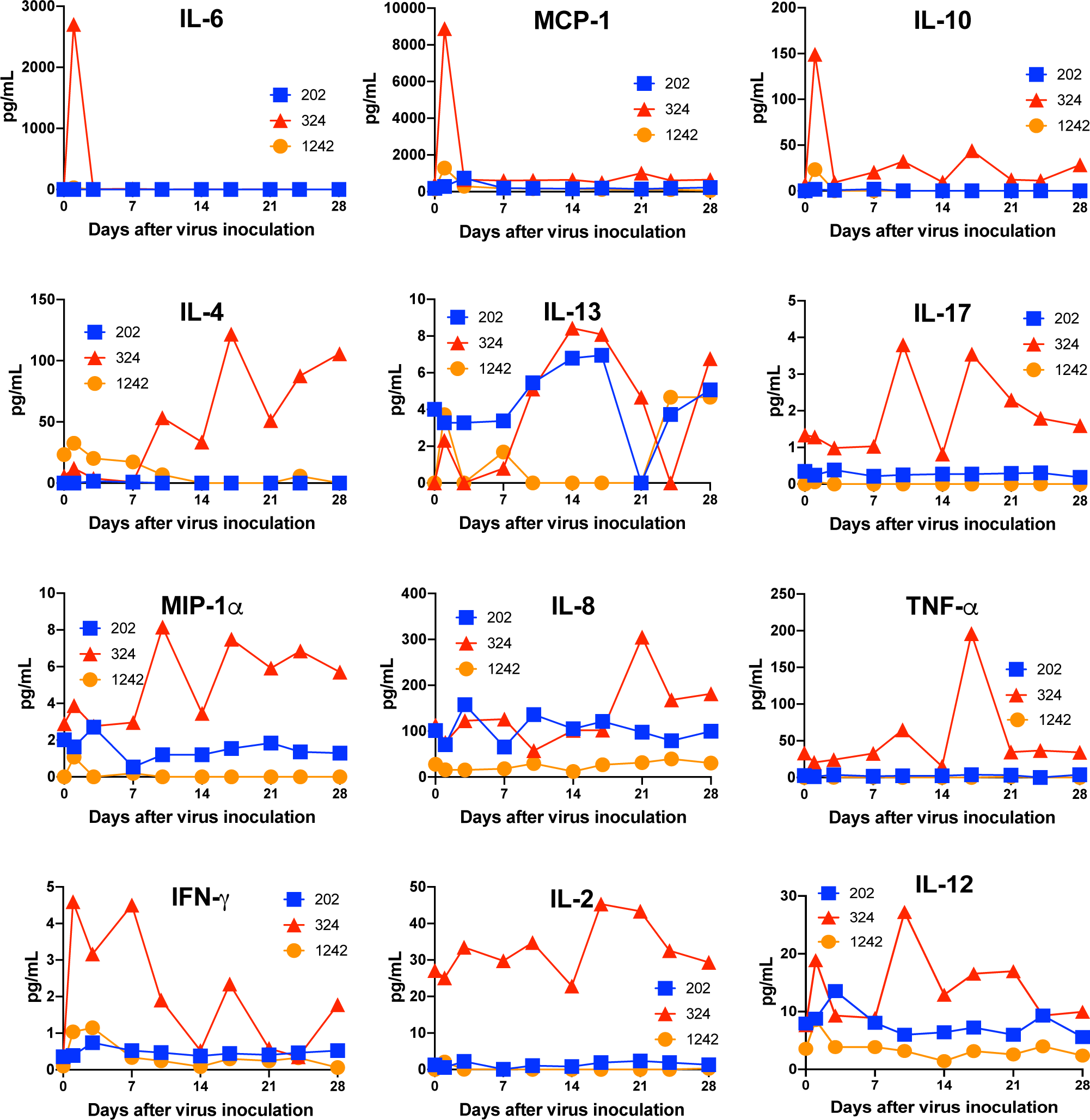
Levels of cytokines and chemokines in plasma of macaques infected with SARS-CoV-2. The concentrations of the indicated cytokines and chemokines in plasma collected on the indicated days from SARS-CoV-2-infected cynomolgus macaques are shown.

The neutralizing antibody against SARS-CoV-2 was detected in one macaque. A low titer of the neutralizing antibody against WK-521 was detected in CE0324F on day 10 after virus inoculation, but no neutralizing antibody was detected in the two other macaques until day 28 (Table 2). An IgG response against SARS-CoV-2 S and N proteins was detected in the plasma of CE0324F and CE1242F after day 14 and day 21, respectively, using an available antibody detection kit for human IgG/IgM (Fig. S3). No antibody response against SARS-CoV-2 was detected in CE0202M, in which the number of BALTs was smaller than those in CE0324F and CE1242F, indicating a relationship between IgG responses and BALT formation.

**Table 2.** Neutralizing antibody in macaques after infection with SARS-CoV-2.

| 50% neutralization titer (log<sub>2</sub>) |  |  |  |
| --- | --- | --- | --- |
| Days after virus inoculation | CE0202M | CE0324F | CE1242F |
| 0 | <<sup>1</sup> | < | < |
| 7 | < | < | < |
| 10 | < | 3.00 | < |
| 14 | < | < | < |
| 17 | < | 3.00 | < |
| 21 | < | 4.00 | < |
| 24 | < | 4.23 | < |
| 28 | < | 5.00 | < |

<sup>1</sup><: 50% neutralization titer was below 3 (2<sup>3</sup>: detection limit).

T lymphocyte responses specific for SARS-CoV-2 N protein were examined. In CE0324F, in which the neutralizing antibody was detected, increases in the number of IL-2 and IFN-γ-producing cells specific for SARS-CoV-2 N protein were seen on day 14 after virus inoculation (Fig. 7). In CE0202M, increases in the number of IFN-γ-producing cells and IL-2-producing cells were detected on day 14 and day 21, respectively. In CE1242F, increases in the number of IFN-γ-producing cells and IL-2-producing cells were detected on day 7 and day 14, respectively. In the two latter monkeys, the increase in the number of IL-2-producing cells followed that of IFN-γ-producing cells and the stimulation indices of IFN-γ-producing cells were higher than those of IL-2-producing cells.

**Fig. 7.**
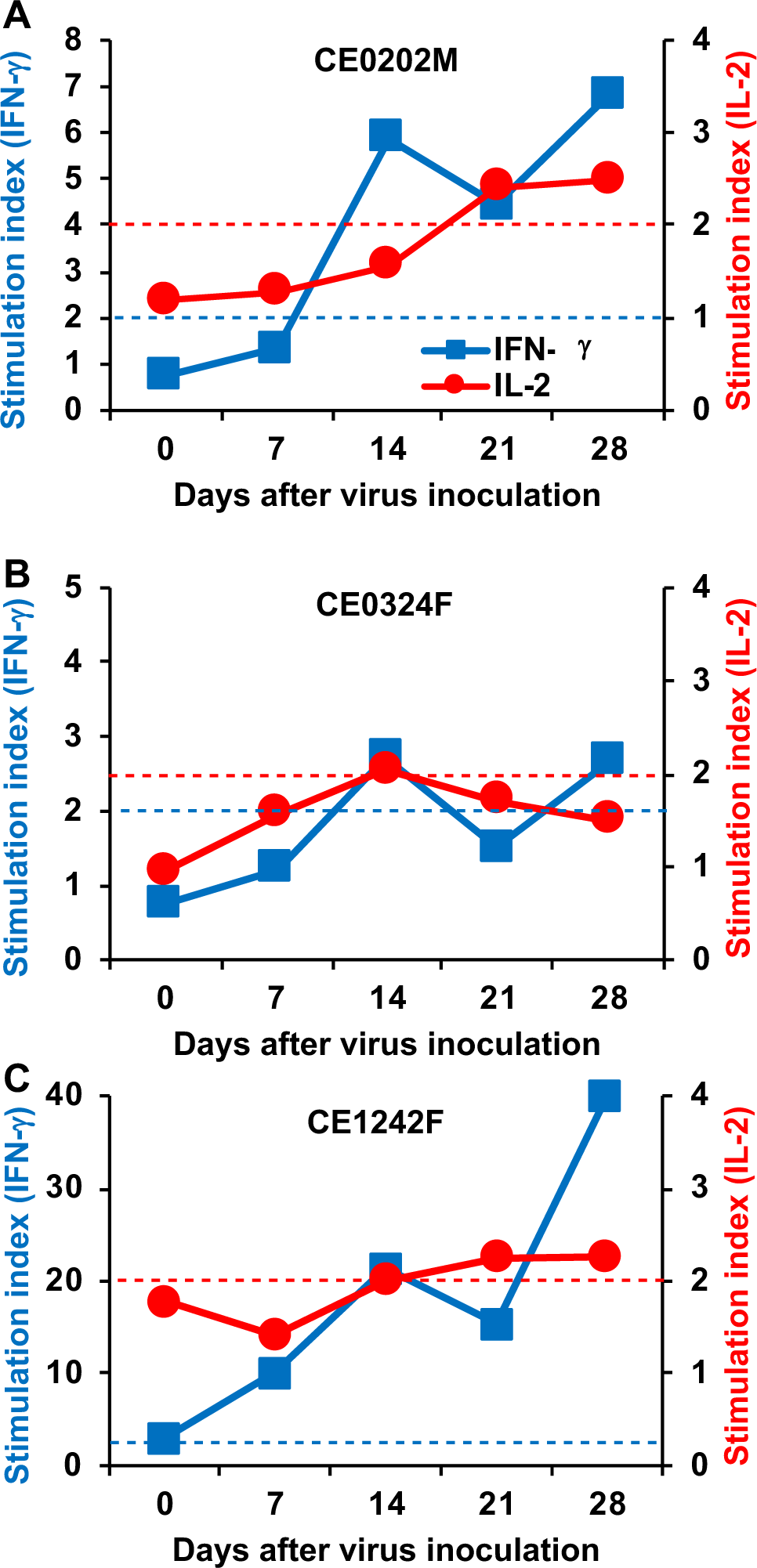
Cytokine responses of T lymphocytes specific for SARS-CoV-2 N protein. Peripheral blood cells collected from macaques (A) CE0202M, (B) CE0324F, and (C) CE1242F on the indicated days were cultured for 24 h with peptides derived from SARS-CoV-2 N protein. The numbers of IFN-γ-positive and IL-2-positive cells were counted using an ELISPOT analyzer. Peptide-specific responses were indicated as a stimulation index = number of spots in the culture with peptides/number of spots in the culture without peptides. Left y-axis: stimulation index of IFN-γ, right y-axis: stimulation index of IL-2. Blue and red dotted lines indicate stimulation indexes 2 in IFN-γ and IL-2, respectively. Stimulation indexes higher than 2 are thought as responses specific for peptide antigens.

## Discussion

We revealed the pathogenicity of SARS-CoV-2 in cynomolgus macaques to establish an animal model for the development of vaccines and antiviral drugs. SARS-CoV-2 was detected predominantly in the nasal cavity and oral cavity of cynomolgus macaques for one week, and infection with SARS-CoV-2 caused a body temperature rise in all of the cynomolgus macaques examined. However, radiographic viral pneumonia was observed in one third of macaques, in which the neutralizing antibody against SARS-CoV-2 was detected. Histologically, pneumonia in the repairing phase and/or bronchopneumonia were observed in all of the macaques and a thrombus in a pulmonary blood vessel was detected [24]. Mild clinical signs of disease and virus propagation in the respiratory tissue of cynomolgus macaques in the present study are consistent with the results of previous studies using cynomolgus macaques and rhesus macaques [11, 12] and with the results of a clinical study showing that approximately 80% of human patients with COVID-19 did not show critical disease [27].

No neutralizing antibody against SARS-CoV-2 was detected 28 days after virus inoculation in two macaques in which body temperature increased and the virus propagated. On the other hand, antibody responses specific for SARS-CoV-2 antigens were detected in most of the macaques on day 14 after infection in previous studies [11, 12]. Our results are consistent with results of studies showing delayed IgM and IgG responses in human patients with COVID-19 [28–30]. The results of the present study indicate that delayed antibody responses would occur in patients without severe symptoms and complications. The results also suggested that the lack of an antibody against SARS-CoV-2 does not necessarily mean evidence of uninfected individuals.

We also examined T-lymphocyte responses specific for SARS-CoV-2 N protein peptides. In macaque CE0324F, in which the neutralizing antibody was detected, the number of cells producing IFN-γ and IL-2 specific for SARS-CoV-2 N protein was increased 2 weeks after virus inoculation. On the other hand, in CE0202M and CE1242F, in which no neutralizing antibody was detected, an increase in the number of cells producing IL-2 specific for SARS-CoV-2 N protein followed an increase in the number of cells producing IFN-γ. The results indicate that an IFN-γ response with a low IL-2 response is not sufficient for induction of a neutralizing antibody. In addition, the IFN-γ response (Th1 response) that helps so-called ‘cellular immunity’ might contribute to elimination of SARS-CoV-2 without a neutralizing antibody. Another possibility is that the early IFN-γ response in CE1242F was induced by memory Th1 cells, suggesting previous exposure to the other coronaviruses. The effects of immunity against the common cold coronavirus on SARS-CoV-2 infection will be analyzed using the macaque model in the future [31].

The level of amylase in the plasma of CE0324F was temporally increased. It has been reported that blood amylase was released from the injured pancreas in patients with COVID-19 pneumonia [32, 33]. However, we did not detect inflammation or necrosis in the pancreas of macaques infected with SARS-CoV-2 at autopsy in the present study (data not shown). On the other hand, lymphoid infiltration in the submandibular gland of CE0324F might be related to the increase of plasma amylase [34], indicating that plasma amylase was derived from infected salivary glands and that the virus detection in oral swab samples was due to virus propagation in the salivary glands [35].

In the present study, we revealed that SARS-CoV-2 propagated in the respiratory tissues of cynomolgus macaques and that SARS-CoV-2 caused clinical signs of disease, indicating that the cynomolgus macaque model is useful for the evaluation of vaccination and therapies. In addition, delayed and low antibody responses in the macaque model suggest that vaccination against SARS-CoV-2 requires high immunogenicity of vaccines and multiple immunizations in humans.

## Materials and methods

### Ethics statement

This study was carried out in strict accordance with the Guidelines for the Husbandry and Management of Laboratory Animals of the Research Center for Animal Life Science at Shiga University of Medical Science and Standards Relating to the Care and Fundamental Guidelines for Proper Conduct of Animal Experiments and Related Activities in Academic Research Institutions under the jurisdiction of the Ministry of Education, Culture, Sports, Science and Technology, Japan. The protocols were approved by the Shiga University of Medical Science Animal Experiment Committee (permit number: 2020-4-2). Regular veterinary care and monitoring, balanced nutrition and environmental enrichment were provided by the Research Center for Animal Life Science at Shiga University of Medical Science. The macaques were euthanized at the endpoint (28 days after virus inoculation) using ketamine/xylazine followed by intravenous injection of pentobarbital (200 mg/kg). Animals were monitored every day during the study to be clinically scored as shown in Table S2 and to undergo veterinary examinations to help alleviate suffering. Animals would be euthanized if their clinical score reached 15 (a humane endpoint).

### Animals

Fifteen-year-old and ten-year-old female cynomolgus macaques (CE0324F and CE1242F) and a fifteen-year-old male cynomolgus macaque (CE0202M) born at Shiga University of Medical Science were used. All procedures were performed under ketamine and xylazine anesthesia, and all efforts were made to minimize suffering. Food pellets of CMK-2 (CLEA Japan, Inc., Tokyo, Japan) were provided once a day after recovery from anesthesia and drinking water was available *ad libitum*. The animals were singly housed in cages equipped with bars to climb up for environmental enrichment under controlled conditions of humidity (39% - 61%), temperature (24 – 26°C), and light (12-h light/12-h dark cycle, lights on at 8:00 a.m.). Two weeks before virus inoculation, a telemetry probe (M00, Data Sciences International, St. Paul, MN) was implanted in the peritoneal cavity or subcutaneous tissue of each macaque under ketamine/xylazine anesthesia followed by isoflurane inhalation to monitor body temperature. The macaques used in the present study were free from herpes B virus, hepatitis E virus, *Mycobacterium tuberculosis*, *Shigella* spp., *Salmonella* spp., and *Entamoeba histolytica*.

Under ketamine/xylazine anesthesia, two cotton sticks (Eiken Chemical, Ltd., Tokyo, Japan) were used to collect fluid samples from the conjunctivas, nasal cavities, oral cavities and tracheas, and the sticks were subsequently immersed in 1 mL of minimal essential medium (MEM, Nacalai Tesque, Kyoto, Japan) containing 0.1% bovine serum albumin (BSA) and antibiotics. A bronchoscope (MEV-2560; Machida Endoscope Co. Ltd., Tokyo, Japan) and cytology brushes (BC-203D-2006; Olympus Co., Tokyo, Japan) were used to obtain bronchial samples. Samples were collected on indicated days.

Chest X-ray radiographs were taken using the I-PACS system (Konica Minolta Inc., Tokyo, Japan) and PX-20BT mini (Kenko Tokina Corporation, Tokyo, Japan). SpO_2_ was measured with a pulse oximeter (Nellcor^TM^, Medtronic plc, Dublin, Ireland).

### Virus and cells

SARS-CoV-2 JP/TY/WK-521/2020 was used as a challenge strain (GenBank Sequence Accession: LC522975, kindly provided by Drs. Masayuki Saijo and Masaaki Sato, National Institute of Infectious Disease (NIID)) [26]. The virus was propagated twice in NIID and once at the Shiga University of Medical Science using VeroE6 (American Type Culture Collection, Manassas, VA) in the presence of BIOMYC-3 (Biological Industries, Beit Haemek, Israel).

The macaques were challenged with the WK-521 virus (2.2 × 10^6^ TCID_50_) inoculated into the conjunctiva (0.05 mL × 2), nostrils (0.5 mL × 2), oral cavity (0.9 mL), and trachea (5 mL) with pipettes and catheters under ketamine/xylazine anesthesia. Experiments using the virus were performed in the biosafety level 3 facility of the Research Center for Animal Life Science, Shiga University of Medical Science.

VeroE6 cells were grown in MEM supplemented with 10% inactivated fetal bovine serum (Capricorn Scientific GmbH), penicillin (100 units/mL), and streptomycin (100 µg/mL) (Nacalai Tesque). To assess virus replication, serial dilutions of swab samples and tissue homogenate samples (10% w/v) were inoculated onto confluent VeroE6 cells. The VeroE6 cells were cultured in MEM supplemented with 0.1% BSA, penicillin, streptomycin, gentamycin (50 μg/mL) (Fuji Film, Tokyo, Japan), and trypsin (5 μg/mL) (Nacalai Tesque). Cytopathic effects were examined under a microscope 6 days later.

### Histopathological examination

For histopathological examination, tissues obtained at autopsy were immersed in 10% neutral buffered formalin for fixation, embedded in paraffin, and cut to 3-μm-thick sections on glass slides. The sections were stained with hematoxylin and eosin (H & E) and observed under a light microscope.

### Blood cytokine and biochemical analyses

Levels of cytokines/chemokines in macaque plasma were measured using the Milliplex MAP non-human primate cytokine panel and Luminex 200 (Millipore Corp., Billerica, MA) following the manufacturer’s instructions. Blood biochemical analysis and blood cell counts were performed using VetScan VS2 and HM2, respectively (Abaxis, Inc., Union City, CA).

### Virus neutralization assay

Plasma samples were pretreated with a receptor-destroying enzyme (RDEII, Denka Seiken, Tokyo, Japan) at 37°C overnight and then inactivated at 56°C for 1 h. The diluted samples were mixed with 100 TCID50/well of the WK-521 strain for 30 min. Then the mixture was added onto a VeroE6 monolayer in 96-well plates. After 1-h incubation, the cells were cultured in MEM containing 0.1% BSA and 5 μg/mL trypsin. After incubation at 37°C for 6 days, the number of wells with cytopathic effects was counted in quadruplicate culture. Neutralization titers were expressed as the dilution in which cytopathic effects were observed in 50% of the wells.

### Detection of an antibody against SARS-CoV-2

Plasma were collected on indicated days. A COVID-19 IgG/IgM immunodetection kit (Novus Biologicals USA, Centennial, CO) was used for detection of IgG and IgM specific for SARS-CoV-2 S and N proteins. Monkey IgM seemed not to react to the human IgM detection system.

### Detection of cytokine-producing cells in ELISPOT

Peripheral blood mononuclear cells after separation from red blood cells were stored at -80°C until use. Thawed cells (5 × 10^5^/well) were cultured with a peptide pool of SARS-CoV-2 N protein (0.6 nmol/mL) (PepTivator, Miltenyi Biotech, Bergisch Gladbach, Germany) in the presence of anti-CD28 antibody (0.1 µg/mL) overnight in ELISPOT plates coated with anti-IFN-γ and IL-2 antibodies (Cellular Technology Limited, Shaker Heights, OH). The number of cytokine-producing cells was counted according to the manufacturer’s instructions.

## Acknowledgements

This work was supported by grants from the Japan Agency for Medical Research and Development (AMED) under Grant Numbers 19fk0108172, 20nk0101615, and 20fk0108276 and a grant from the Ministry of Education, Culture, Sports, Science and Technology, Japan for a Joint Research Program of the Research Center for Zoonosis Control, Hokkaido University. Misako Nakayama is supported by the Naito Foundation. Cong Thanh Nguyen is supported by the Sato Yo International Scholarship Foundation. We thank Drs. Kazumasa Ogasawara, Michinori Kohara, Fumihiko Yasui, Masashi Shingai, and Marumi Ohno for providing suggestions, Drs. Masayuki Saijo and Masaaki Sato for providing the virus, Drs. Hideaki Tsuchiya, Ikuo Kawamoto, Takahiro Nakagawa, and Iori Itagaki for animal care, and Hideaki Ishida, Naoko Kitagawa, Takako Sasamura, and Chikako Kinoshita for assistance in experiments.

