## Supplementary material for "Neutralizing antibody-dependent and -independent immune responses against SARS-CoV-2 in cynomolgus macaques": SARS1Fig200813Supplement.pdf

Fig. S1

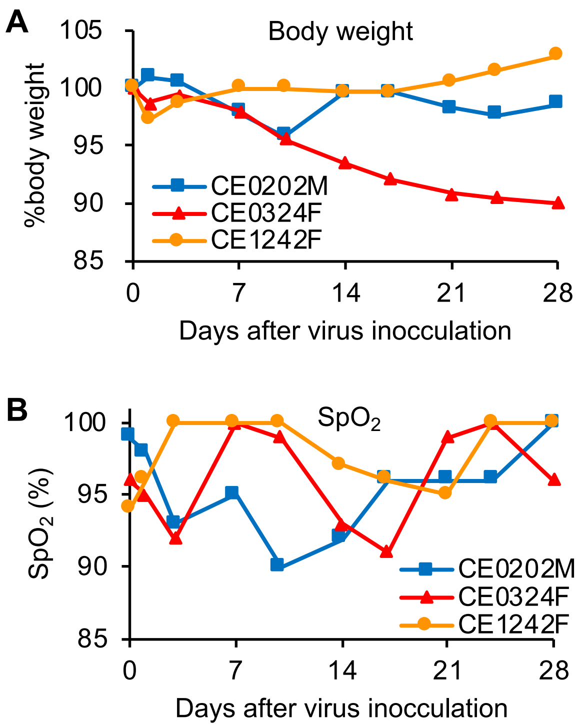


**Figure S1. Body weight and $Sp0_2$ in macaques infected with SARS-CoV-2.**

(A) Body weights and (B) saturation of percutaneous oxygen ($SpO_2$) were measured at sampling under anesthesia.

Fig. S2

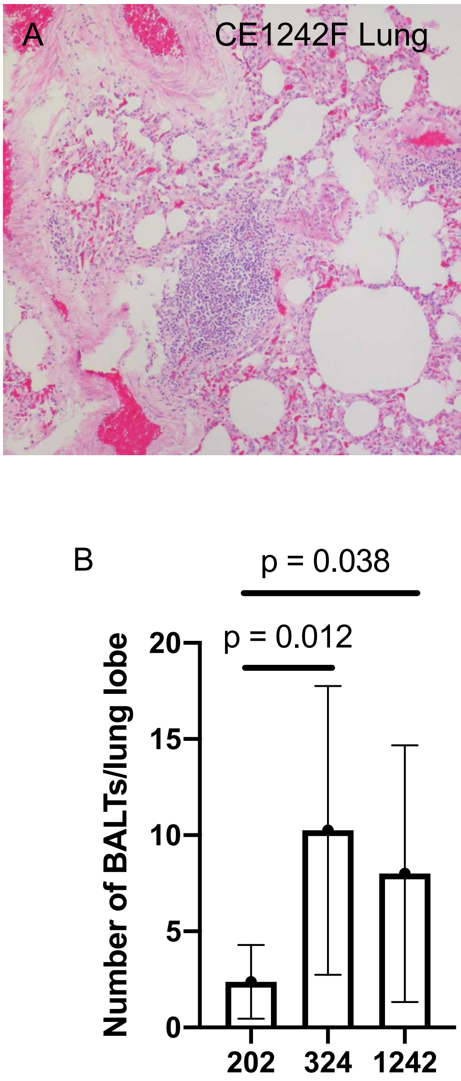


**Figure S2. Number of BALTs in macaques infected with SARS-CoV-2.**
Lung tissues were collected on day 28 after virus inoculation. (A) BALTs in the lung tissue of CE1242F. (B) The number of BALTs in 8 H&E sections of 6 lobes in each macaque was counted by detection of follicle-like structures composed of densely packed lymphocytes, of which the shortest diameter exceeded 0.1 mm, and flanked by the bronchus or vessels. The averages and standard deviations of the number of BALTs in 8 sections are shown. Unpaired t test (2 tailed) was performed.

Fig. S3

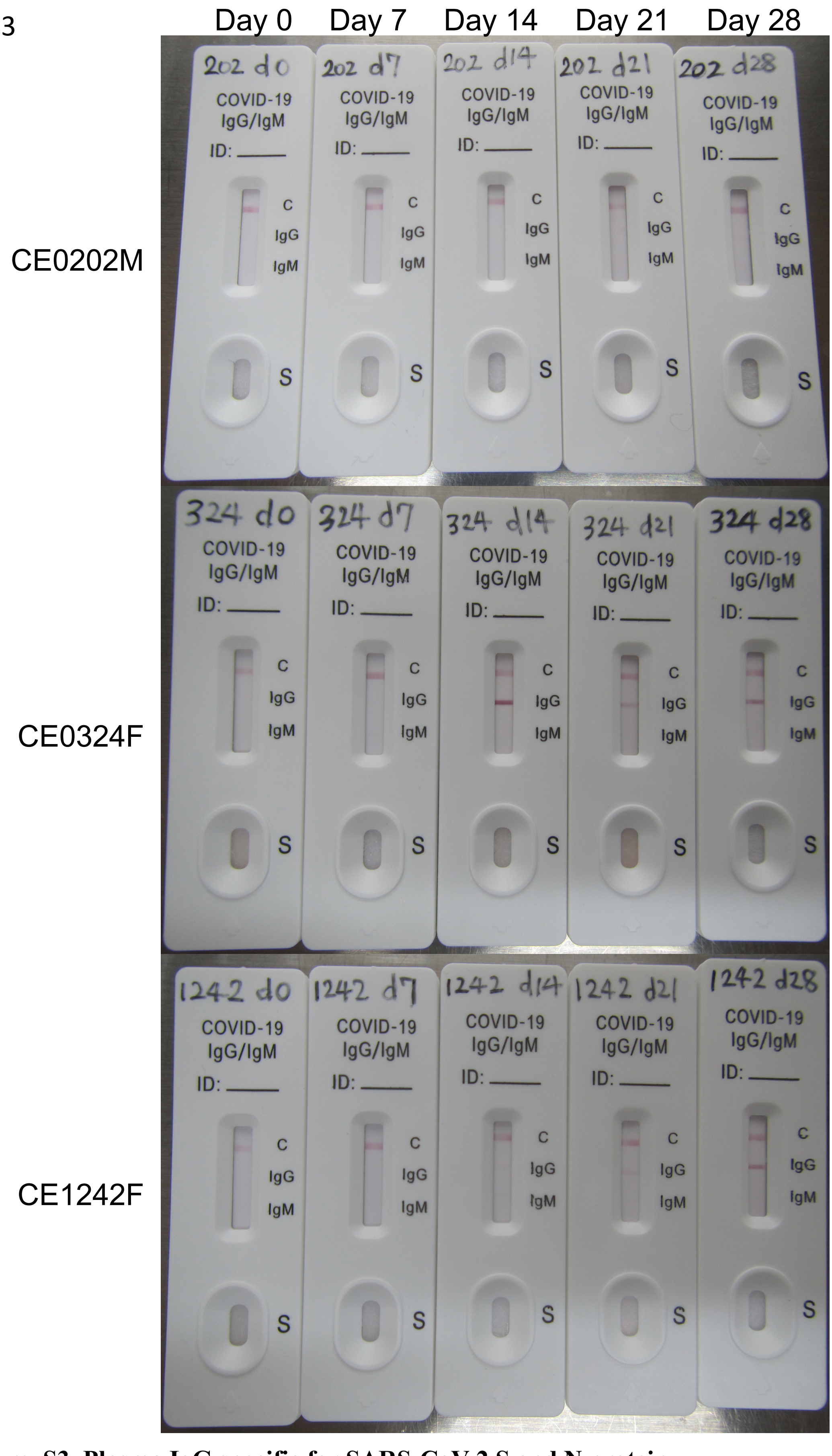


**Figure S3. Plasma IgG specific for SARS-CoV-2 S and N protein**.
Blood samples were collected from macaques day 0, 7, 14, 21 and 28. IgG specific for SARS-CoV-2 S and N protein was detected in CE0324F and CE1242F

**Table S1. Tissues collected at autopsy**

| Respiratory tissue | Brain | Digestive organs | Other organs |
|---|---|---|---|
| Nasal mucosa | Frontal lobe | Liver | Heart |
| Oronasopharynx | Temporal lobe | Jejunum | Spleen |
| Tonsil | Parietal lobe | Ileum | Kidney |
| Trachea | Occipital lobe | Transverse colon | Muscle |
| Bronchus | Olfactory bulb | Descending colon | Uterus |
| Lung, 6 lobes | Cerebellum | Rectum | Ovary |
| Lymph node, mediastinum | Brain stem | | Testis |
| | | | Conjunctiva |
| | | | Eyeball |

The tissues were collected on day 28 to prepare 10% (w/v) homogenates and to titrate for virus detection. No virus was detected in the listed tissues.

**Table S2. Clinical scoring used in the present study**

| Parameter | Degree of parameter | Possible score |
| --- | --- | --- |
| Fever | Normal (< 39°C) | 0 |
| | Elevated temperature (39-40°C) | 3 |
| | High temperature (> 40°C) | 5 |
| Posture | Piloerection of body hair | 1 |
| | Decreased activity, decreasing normal behavior/Occasionally lying down, huddled, active when people in room | 2 |
| | Huddled on camera, active when people in room/Lying down, getting up when approached, using cage for support | 3 |
| | Huddled when people in room, shaking, toes and hands clenched/Lying down, not getting up when approached or prompted | 5 |
| Respiration | Increased or decreased; mild cough and clear nasal discharge | 3 |
| | Labored breathing through mouth; severe cough and severe nasal discharge | 5 |
| Appetite | Slightly decreased | 1 |
| | Decreased | 2 |
| | Severely decreased | 5 |
| Skin | Flushed appearance | 2 |
| | Visible rash | 2 |
| | Bleeding | 5 |

Animals were monitored every day during the study to be clinically scored.
Animals were euthanized if their clinical scores reached 15 (a humane endpoint).
